# MSCs from syndecan-3 null mice exhibit enhanced adhesion to collagen type I, hyperactivation of the AKT pathway and increased efficacy in inflammatory arthritis

**DOI:** 10.1101/2020.06.12.148221

**Authors:** Fiona K Jones, Andrei Stefan, Alasdair G Kay, Mairead Hyland, Rebecca Morgan, Nicholas R Forsyth, Addolorata Pisconti, Oksana Kehoe

## Abstract

Rheumatoid arthritis (RA) is a debilitating and painful inflammatory autoimmune disease characterised by the accumulation of leukocytes in the synovium, cartilage destruction and bone erosion. The immunomodulatory effects of bone marrow derived mesenchymal stem cells (MSCs) has been widely studied and the recent observations that syndecan-3 (SDC3) is selectively pro-inflammatory in the joint led us to hypothesise that SDC3 might play an important role in MSC biology. MSCs isolated from bone marrow of wild type and Sdc3^−/−^ mice were used to assess immunophenotype, differentiation, adhesion and migration properties and cell signalling pathways. While both cell types show similar differentiation potential and forward scatter values, the cell complexity in wild type MSCs was significantly higher than in Sdc3^−/−^ cells and was accompanied by lower spread surface area. Moreover, Sdc3^−/−^ MSCs adhered more rapidly to collagen type I and showed a dramatic increase in AKT phosphorylation, accompanied by a decrease in ERK1/2 phosphorylation compared with control cells. In a mouse model of antigen-induced inflammatory arthritis, intraarticular injection of Sdc3^−/−^ MSCs yielded enhanced recovery compared to injection of wild type MSCs. In conclusion, our data suggest that syndecan-3 regulates MSC adhesion and efficacy in inflammatory arthritis, likely via induction of the AKT pathway.

## 2 Introduction

Syndecans are transmembrane heparan sulphate proteoglycans (HSPG) that consist of a core protein with heparan sulphate glycosaminoglycan chains covalently attached to the ectodomain. They form part of the glycocalyx, which comprises a network of membrane-bound proteoglycans and glycoproteins on the surface of many different cell types^1^. There are four mammalian syndecans, designated syndecan-1 (SDC1), -2, -3, and -4, which have protein cores with characteristic structural domains^2^. The variable ectodomain, which is exposed to the extracellular environment, contains two or more heparan sulphate chains and in some cases also chondroitin sulphate chains. Through their ectodomain syndecans interact with a variety of extracellular and membrane proteins thus modulation many signalling pathways and biological processes including adhesion, proliferation, differentiation and inflammation^3,4^. The hydrophobic transmembrane domain is followed by a short intracellular domain containing two domains highly conserved across all syndecans and a small highly variable domain which is unique across all four syndecans. The syndecan intracellular domain binds to cytoskeletal proteins, scaffold proteins and intracellular kinases involved in signal transduction and gene expression regulation, thus transducing signals collected by the extra cellular domain to both the cytoplasm and the nucleus^5,6^.

Syndecan-3 (SDC3) has the largest core protein of all four mammalian syndecans and can harbour both heparan sulphate and chondroitin sulphate chains^7^. SDC3 is expressed in several cell types and is involved in the regulation of growth factor signalling^8,9^, Notch and BMP signalling^10,11^, adhesion and migration^12,13^, proliferation^10,14^ and differentiation^10,15^.

Inflammation is a central feature of rheumatoid arthritis that affects around 1% of the population and can result in disability and morbidity. Chemokines are involved in stimulating the infiltration of leukocytes into inflamed tissue and the expression of a CXCL8 binding site on endothelial SDC3 in human RA suggests a role for this HSPG in inflammatory disease^16^. We demonstrated that SDC3 plays a dual role in inflammation depending on the tissue and vascular bed. In the joint it is pro-inflammatory, since its deletion leads to reduced leukocyte recruitment and the severity of arthritis. However, in the skin and cremaster SDC3 plays an anti-inflammatory role, since its deletion leads to enhanced leukocyte interaction with the endothelium and recruitment^17^. Furthermore, an article recently published by Eustace *et al.* characterises the role of soluble SDC3 in inflammation using in vitro and in vivo models^18^. The authors show that soluble SDC3 binds chemokines, reduces leukocyte migration in vitro and ameliorates disease severity in mouse models of arthritis. The observed anti-inflammatory mechanisms of shed SDC3 appear to be associated with competitive binding of SDC3 to chemokines, which limit their availability to recruit inflammatory cells^18^.

Mesenchymal stem cells (MSCs) play a major role in the maintenance and regeneration of adult tissues subjected to physiological turnover or following injury^19^. MSCs exert immunomodulatory functions, including inhibition of T cell proliferation, interference with B cell function and dendritic cell maturation and promotion of anti-inflammatory macrophage-mediated responses^20,21,22^. We previously demonstrated that MSCs reduce inflammation, joint swelling and cartilage destruction in a murine model of antigen-induced arthritis (AIA)^17^ and hypothesise that SDC3 might play an important role in MSCs biology. A variant of SDC3 is expressed on the surface of human bone marrow stromal cells where it may be responsible for creating a stromal ‘niche’ for the maintenance and development of hemopoietic and progenitor cells^23^. Here, we show a role for SDC3 deletion on various murine MSC phenotypes, both in vitro and in vivo. MSCs isolated from syndecan-3 null mice show enhanced adhesion to collagen type I and hyperactivation of the AKT pathway, which are associated with increased efficacy of MSC treatment in a mouse model of AIA. Thus, our data further support the idea that pharmacologic targeting of SDC3 might constitute a potentially valuable therapeutic strategy for inflammatory arthritis.

## 3 Results

### 3.1 Characterisation of murine bone marrow MSCs

Wild type and SDC3 murine MSCs were successfully differentiated towards adipogenic, osteogenic and chondrogenic lineages after 21 days in culture with the relevant differentiation media. The differentiation potential was similar for both wild type and *Sdc3*^−/−^ mMSCs (Fig. 1a). A colony-forming units assay was used to assess the proliferative capacity of the cells being expanded in culture, and the results showed both genotypes retained high proliferation rate in culture (Fig. 1b). The mean population doubling time (PDT) value for *Sdc3*^−/−^ MSCs was lower than that for wild type cells but was not statistically significant (Fig. 1c). Immunophenotypic analysis indicated similar surface marker expression pattern for both wild type and *Sdc3*^−/−^ MSCs. Both sets of cells were negative for haematopoietic markers CD11b and CD45 and for the endothelial cell markers CD31, and positive for the mesenchymal markers CD44 and Sca-1. Expression of the general progenitor cell marker CD34 and of the vascular progenitor marker CD105 was highly variable in MSCs isolated from both genotypes, with a general trend of both markers being more commonly expressed in MSCs from *Sdc3*^−/−^mice compared with MSCs from wild type mice (Fig. 1d and e). Flow cytometry showed similar mean values for cell size (FSC, forward scatter) for both cell types (wild type: 112146.53±6324.64; *Sdc3*^−/−^: 102259.3533±3460.25) (Fig. 1f). However, the spread cell surface area of adherent *Sdc3*^−/−^ MSCs was considerably lower when compared with wild type (Fig. 1g). Similarly, wild type MSCs showed significantly higher cellular complexity, measured by side scatter (SSC) values then the *Sdc3^−/−^* MSCs (Fig. 1h).

**Figure 1.**
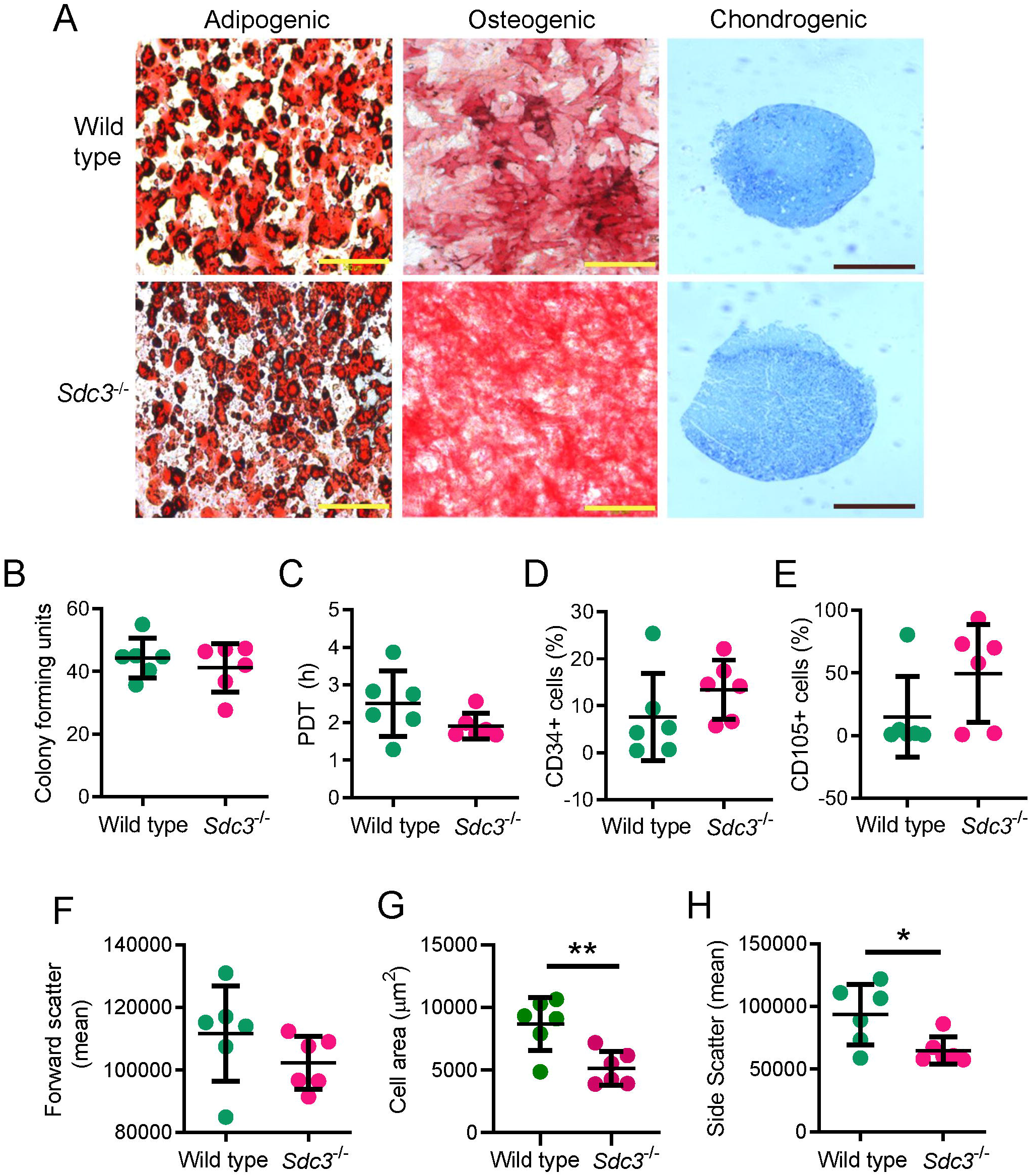
Characterisation of murine bone marrow MSCs. **(a)** Wild type and *Sdc3*^−/−^ MSCs show a similar trilineage differentiation potential *in vitro*. Yellow scale bar represents 200 μm; black scale bar represents 500 μm. The proliferative capability of MSCs were measured by **(b)** fibroblast colony-forming units and **(c)** average population doubling time (PDT). Dot plots displaying the percentage of cells expressing CD34 **(d)** or CD105 **(e)** as measured by flow cytometry. **(f)** The geometric mean of cell size was determined by forward scatter. **(g)** The morphometry, or cell spread, of cells were analysed using ImageJ. **(h)** The internal complexity of cells was measured by side scatter and the geometric means are shown. MSCs were isolated from wild type (n=6) and *Sdc3*^−/−^ (n=6) mice. Data presented as mean ± standard deviation unless otherwise stated; * = p< 0.05; ** = p< 0.01.

### 3.2 Syndecan-3 loss improves adhesion on collagen without affecting MSC migration

Crosstalk between extracellular matrix (ECM) components and MSCs is essential for tissue formation and maintenance. Syndecan-3 participates directly in cell adhesion the ECM as a collagen receptor, and might act as an integrin co-receptor by analogy to its close homolog SDC1^13,24–27^. Thus, we tested wild type and *Sdc3^−/−^* MSC adhesion to various ECM substrates. Without ECM coating, approximately 65% of wild type and *Sdc3*^−/−^ cells were attached to tissue culture treated plates within 1 hour of incubation (Fig. 2a), however, when seeded onto collagen-coated tissue culture plates, *Sdc3*^−/−^ MSCs adhered more quickly than wild type MSCs (p=0.014) (Fig. 2b). Similarly, fibronectin improved *Sdc3*^−/−^ MSC adhesion by ~19% compared with wild type MSCs, although this difference was not statistically significant (Fig. 2c). Interestingly, laminin coating appeared to yield the opposite effect of collagen and fibronectin coating, although the difference in adhesion to laminin between wild type and *Sdc3*^−/−^ MSCs was not statistically significant (Fig. 2d).

**Figure 2.**
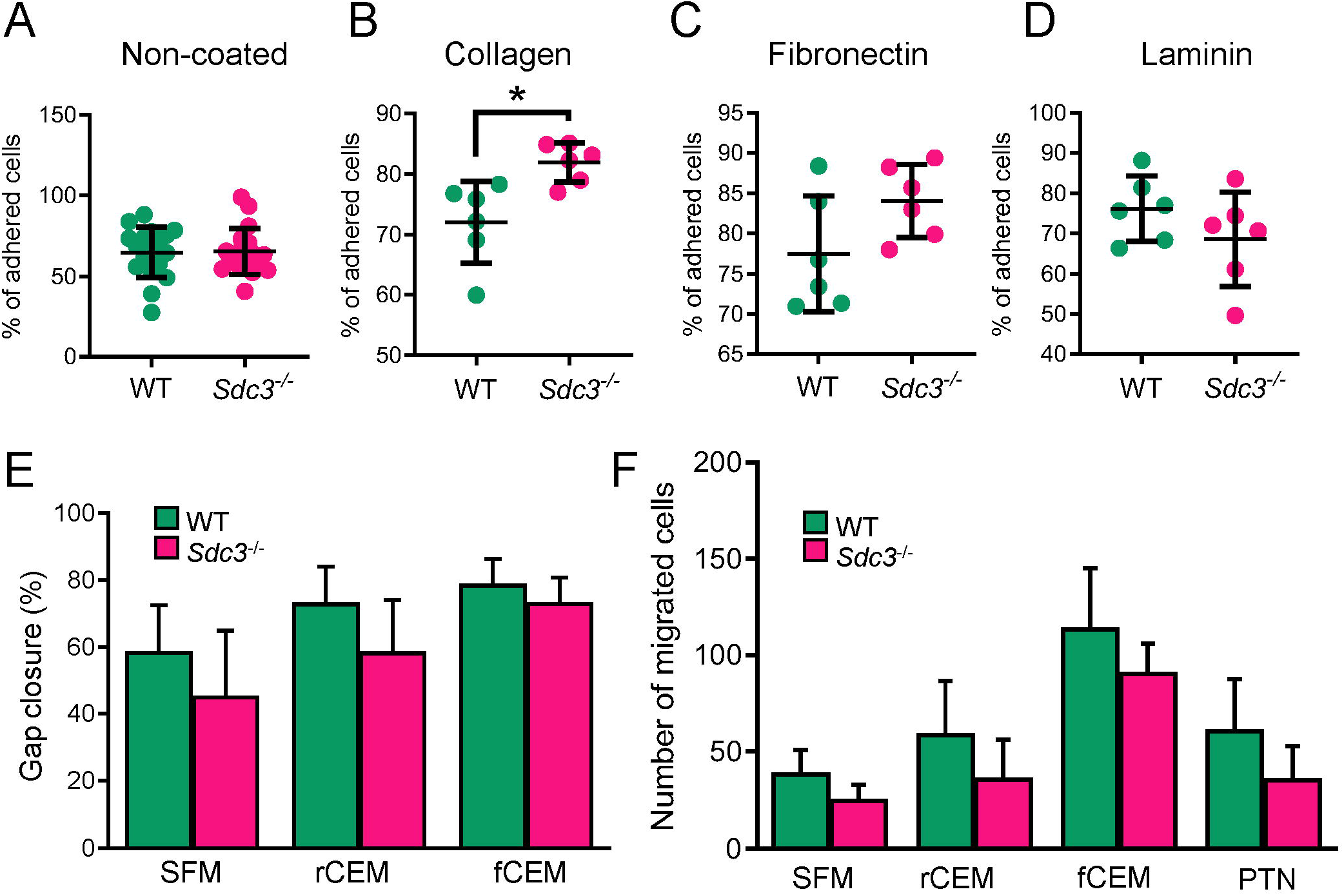
Adhesion to collagen is increased in *Sdc3^−/−^* MSCs. MSCs were isolated from wild type (n=6) and *Sdc3*^−/−^ (n=6) mice and seeded onto several different substrates in complete expansion medium (CEM). After 60 min the average percentage of wild type (WT) and *Sdc3*^−/−^ MSCs adhered to each substrate was calculated: **(a)** Non-coated, **(b)** Collagen type I, **(c)** Fibronectin and **(d)** Laminin. **(e)** A scratch assay was performed on MSCs cultured in serum free medium (SFM), reduced serum content medium (rCEM) or full serum content medium (fCEM) and the average percentage of gap closure is shown. **(f)** Cell migration towards specific chemoattractants was measured using a transwell assay. The number of migrated cells was calculated after 15 h of incubation to each chemoattractant. PTN, pleiotrophin. Data presented as mean ± standard deviation; * P< 0.05.

We further analysed the effect of syndecan-3 deletion on cell migration by the scratch and transwell assays. When cells were cultured in serum-free medium (SFM) or complete expansion medium (CEM) with either full serum content (FBS and HS, fCEM) or reduced serum content (CEM with 1% FBS and 1% HS, rCEM) in a scratch assay, wild type MSCs migrated slightly faster than *Sdc3^−/−^* MSCs but differences were not significant (Fig. 2e). To further investigate a potential role for SDC3 in MSC migration, we tested cell migration towards specific chemoattractants using a transwell assay. In the absence of chemoattractants, *Sdc3*^−/−^ MSCs showed a tendency to migrate less than wild type MSCs, and although serum addition to the lower chamber increased MSC migration in a dose-dependent manner, the difference in migration between wild type and *Sdc3*^−/−^ MSCs remained constant and non-statistically significant (Fig. 2f). Pleiotrophin is a major soluble ligand for SDC3 and acts as a chemoattractant for osteoblasts and neural progenitors in a SDC3-dependent manner^12^. When pleiotrophin was added to low serum concentrations in the lower chamber of transwells it failed to enhance MSC migration (Fig. 2f). All these data together suggest that the slight migration defect observed in *Sdc3^−/−^* MSCs is due to an intrinsic property of *Sdc3*^−/−^ MSCs rather than to a role for SDC3 in sensing external chemoattracting cues.

### 3.3 Syndecan-3 regulates ERK and AKT signalling pathways in MSCs

The results shown so far suggest a role for SDC3 in the regulation of MSC adhesion via one or more collagen receptors. Typical collagen receptors are integrins, which signal via several kinase-regulated pathways, and some tyrosine kinase receptors^28,29^. Previous work demonstrated that *Sdc3*^−/−^ satellite cell-derived myoblasts had a global increase in phosphorylated-tyrosine (pY), suggesting SDC3 is involved in regulating numerous signalling pathways simultaneously. Therefore, we first measured the levels of pY by western blot in wild type and *Sdc3*^−/−^ MSCs but found no difference (Fig. 3a and b) and found no differences between the two genotypes.

**Figure 3.**
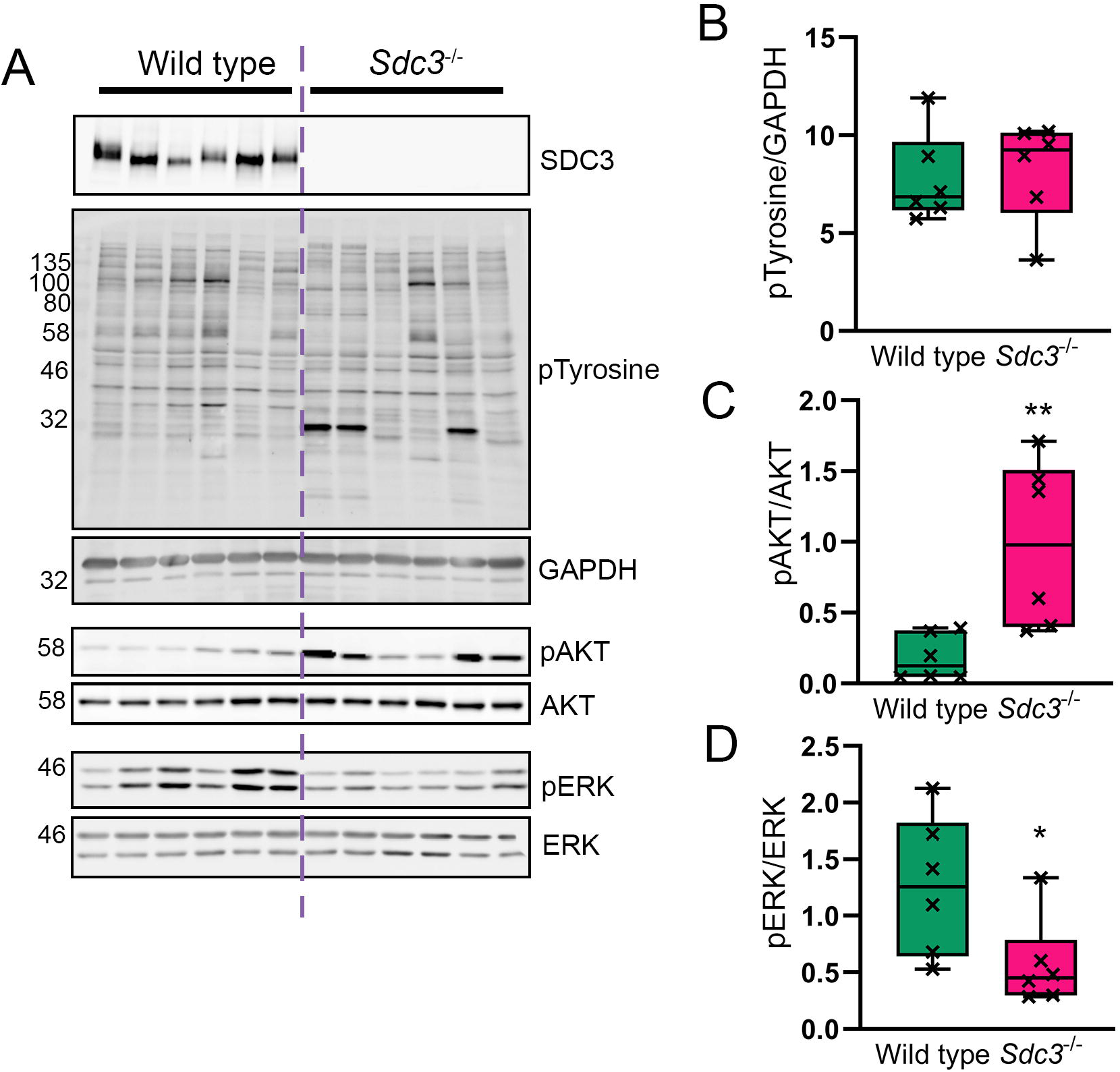
Loss of SCD3 in MSCs causes increased ATK phosphorylation and a reduction in ERK phosphorylation. MSCs were isolated from wild type (n=6) and *Sdc3*^−/−^ (n=6) mice, cultured in full growth medium, and lysed ready for western blotting. **(a)** representative images of blots used for quantification. SDC3 knockout was confirmed for *Sdc3*^−/−^ MSCs in each sample **(b)** Total phosphorylated tyrosine normalised to GAPDH. **(c)** Phosphorylated AKT^Ser473^ was normalised to total AKT. (D) Phosphorylated ERK^Thr202/Tyr204^ was normalised to total ERK. Quantification was performed using ImageJ. Data displayed as mean ± standard deviation; * = p< 0.05; ** = p< 0.01.

Common kinase pathways downstream several adhesion complexes are the RAS/ERK pathways and the PI3K/AKT pathway^30^. Thus, we investigated changes in phosphorylated AKT and ERK1/2. Here we found that the levels of pAKT^S473^ were significantly increased in *Sdc3^−/−^* MSCs compared to wild type MSCs (Figure 3c). In contrast, the levels of pERK1/2 significantly decreased when SDC3 was lost from MSCs (Figure 3d), suggesting a role for SDC3 in inhibiting AKT signalling while promoting ERK1/2 signalling in MSCs adhering to collagen.

### 3.4 Syndecan-3 loss in MSCs improves arthritic symptoms in AIA

The anti-inflammatory capability of MSCs from wild type and *Sdc3*^−/−^ mice were tested in the AIA mouse model of inflammatory arthritis. This model shares many histopathological and clinical features seen in human RA^31^. MSC treatments (wild type and *Sdc3*^−/−^) reduced joint swelling as a measure of joint inflammation compared to serum-free medium control at days 2 (p<0.05) and 3 (p<0.05) post arthritis induction (Fig. 4a). Significant reductions were also recorded in exudates in the joint cavity and cartilage loss at day 3 following MSC treatment and in overall arthritis index at day 3 post-arthritis induction (Fig. 4b). At day 7 post-arthritis induction, the histological scores were reduced but not significantly (Supplementary Fig. 1). There were no significant differences in the histological parameters between knees treated with MSCs from wild type mice and knees treated with MSCs from *Sdc3*^−/−^ mice.

**Figure 4.**
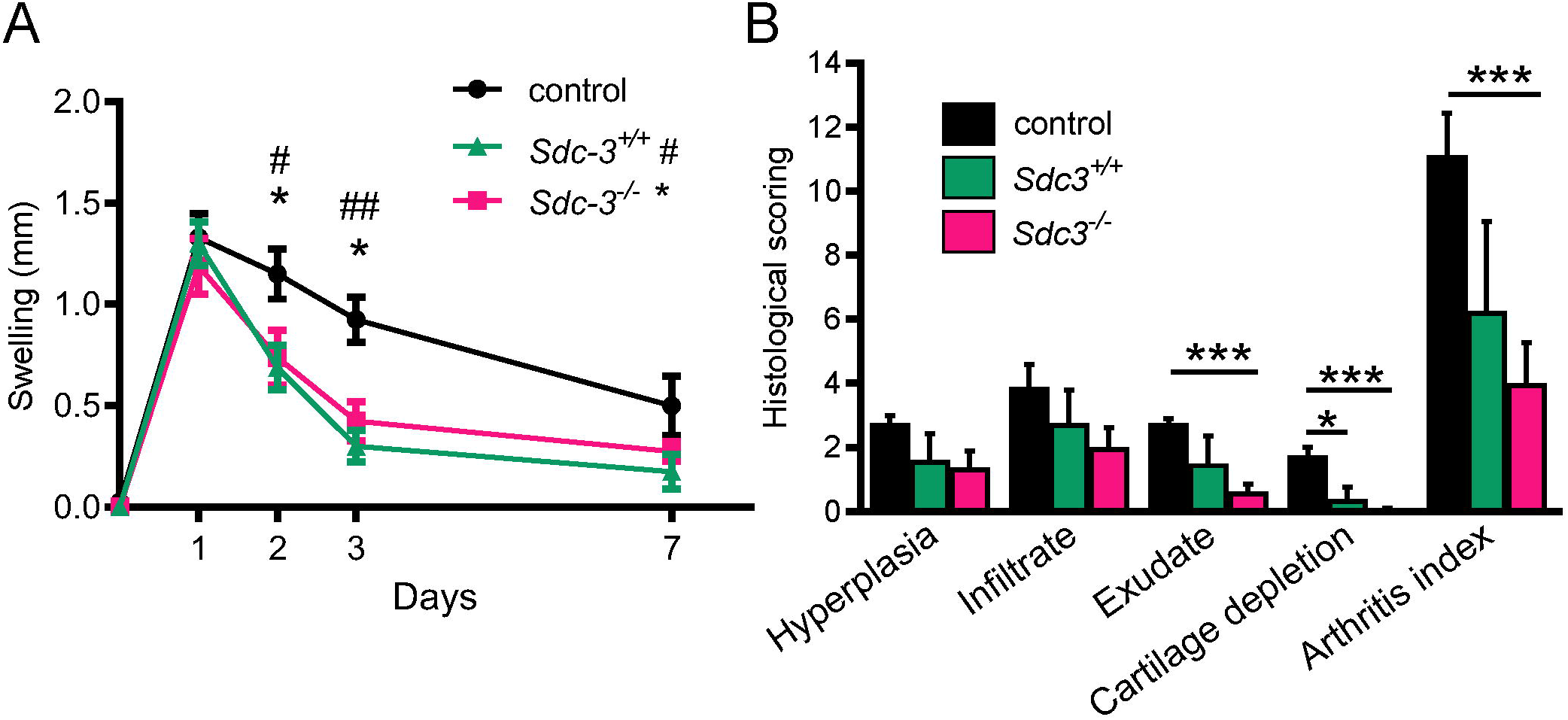
Effects of intra-articular injections of MSCs in AIA. **(a)** Knee diameter (mm) was used as an index of swelling (joint inflammation) and measured at days 1,2,3 and 7 post arthritis induction. Significant reductions are seen following *Sdc3*^−/−^ MSCs and *Sdc3^+/+^* MSCs injection in AIA mice. Data are means ± standard error of the mean (SEM) for right knee after subtraction of left knee control. * p<0.05 comparing *Sdc3*^−/−^ MSCs and control mice at the time points indicated; # p<0.05 and ## p<0.01 comparing *Sdc3^+/+^* MSCs and control mice at the time points indicated. **(b)** The histological scoring for 3 days post arthritis induction. The arthritis index is the sum of all observations. Data are means ± SEM, n=4 mice in each group; *p<0.05 and ***p<0.001.

## 4 Discussion

In this work we describe a role for the transmembrane proteoglycan SDC3 in MSC biology, with a particular focus on its role as an adhesion molecule and its involvement in MSC-mediated promotion of cartilage repair in a model of inflammatory arthritis.

Phenotypic characterisation of MSCs isolated from *Sdc3*^−/−^ mice showed that the overall effect of SDC3 loss in mice does not produce dramatic effects on MSCs as *Sdc3*^−/−^ retained multilineage potential *in vitro* and similar proliferation rates as wild type MSCs. Immunophenotyping analysis also revealed similar surface marker expression pattern for both wild type and *Sdc3*^−/−^ MSCs. However, endoglin (CD105), a transmembrane glycoprotein that acts as an accessory to the transforming growth factor β (TGFβ) receptor system^32^, was more commonly expressed in MSCs from *Sdc3*^−/−^ mice compared with MSCs from wild type mice. Endoglin is an MSC marker and it is expressed in progenitor cells involved in vascular remodelling in animal models of myocardial infarction and rheumatic diseases^33,34^. It has been shown that endoglin promotes angiogenesis in cell- and animal-based models of retinal neovascularisation and treatment with anti-CD105 antibody inhibits neovascularisation in oxygen-induced retinopathy model^35^. Similarly, it has been shown that SDC3 inhibits angiogenic factor *in vitro*^36^ and that regenerating muscles of *Sdc3*^−/−^ mice show enhanced angiogenesis^15^. Thus, it is possible that the increase in CD105 expression in response to SDC3 loss might be linked to the SDC3-mediated regulation of angiogenesis that has been observed both *in vitro* and *in vivo*.

Obtaining a sufficient number of cells to initiate cell therapy is a limiting factor during therapeutic applications. Cells must be expanded *in vitro* prior to *in vivo* administration and cell adhesion is necessary for MSC expansion and further therapeutic use^37^. Our results showed a role for SDC3 in the regulation of MSC adhesion via one or more collagen receptors. Previous work demonstrated the cooperation of α2β1 integrin with SDC1 during adhesion to collagen^38^, and cooperation of α2β1 integrin and α6β4 integrin with syndecans during adhesion to laminin^39,40^. A recent study has further confirmed that collagen promotes cell proliferation, cell survival under stress and a stronger cell adhesion to the cell culture surface^37^. Since the attachment of MSC to collagen is mostly β1 integrin-mediated^41^, it is likely that SDC3 regulates MSC adhesion to collagen via β1 integrin.

Cell migration is an essential biological process involved many aspects of health and disease, including development, tissue regeneration, inflammation, and cancer progression. In this study we used two *in vitro* assays to study MSC migration, With the scratch assay, we studied the effects of cell–matrix and cell–cell interactions on cell migration; with the transwell assay we studied the ability of cells to directionally respond to chemoattractants^42^. Our results showed that wild type MSCs migrated slightly faster than *Sdc3*^−/−^ MSCs in both assays, suggesting a role for SDC3 in the regulation of cell motility. Whether SDC3-mediated regulation of MSC migration is directly linked to SDC3-mediated regulation of adhesion or via other pathways, remains to be established. In other cell types SDC3 regulates migration by functioning as a ligand for the soluble factor heparin-binding growth-associated molecule (HB-GAM) also known as pleiotrophin^43,44^. However, pleiotrophin failed to enhance migration of both wild type and *Sdc3*^−/−^ MSCs in our transwell assays, suggesting that, at least under our experimental conditions, SDC3-mediated regulation of MSC migration is not pleiotrophin-dependent. Moreover, since we observed the same slight difference in migration between wild type and *Sdc3*^−/−^ MSCs regardless of the chemoattractant used, we conclude that such migration defect is due to an intrinsic property of *Sdc3*^−/−^ MSCs rather than to a role for SDC3 in sensing external chemoattracting cues.

As discussed above, our data suggest that SDC3 regulates integrin-mediated MSC adhesion to collagen. Indeed, *Sdc3*^−/−^ mMSCs cultured on collagen showed a dramatic increase in AKT phosphorylation, which was accompanied by a decrease in ERK1/2 phosphorylation compared with wild type controls. The adhesion and signalling phenotypes shown by *Sdc3^−/−^* MSCs are mirrored by the phenotypes reported for loss of α2 and α11 integrin, both collagen I receptors, in human MSCs, which leads to decreased AKT phosphorylation, MSC adhesion to collagen I and cell spreading^45^. Thus, our data strongly suggest a role for SDC3 in MSC adhesion and signalling by regulating integrin-collagen interaction.

The immunomodulatory effects of bone marrow derived mesenchymal stem cells (MSCs) has been widely studied and this current study has attempted to characterise the role that SDC3 plays in MSC biology. Deletion of SDC3 led to improvement of clinical scores, leukocyte recruitment and cartilage damage in a murine model of antigen-induced arthritis^17^ suggesting that SDC3 contributes to the clinical manifestation of the disease. In the current work experiments, MSC treatments (wild type and *Sdc3*^−/−^) significantly reduced exudates in the joint cavity and cartilage loss at day 3 following MSC treatment and the overall arthritis index at day 3 post-arthritis induction. *Sdc3^−/−^* MSCs were more effective than wild type cells at reducing the overall disease severity. SDC3 deletion enhances the efficacy of MSC treatment in an AIA model of inflammatory arthritis. Our study indicates important role for SDC3 in MSC biology which requires further investigation.

## 5 Materials and methods

### 5.1 Mice

Wild type mice, C57Bl/6 background, for MSC isolation and for the inflammatory arthritis model were obtained from Envigo, UK and housed in a pathogen-free facility at Liverpool John Moores University, in accordance with the Animals (Scientific Procedures) Act 1986 and the EU Directive 2010/63/EU, and after local ethical review and approval by the Animal Welfare and Ethical Review Body. Syndecan-3 null (*Sdc3^−/−^*) mice, sharing the same C57Bl/6 background as wild type mice (Reizes O et al, 2001), were donated by Dr Rauvala, University of Helsinki and housed in a pathogen-free facility at the University of Liverpool, in accordance with the Animals (Scientific Procedures) Act 1986 and the EU Directive 2010/63/EU, and after local ethical review and approval by the Animal Welfare and Ethical Review Body.

### 5.2 Isolation, expansion, and characterisation of MSCs

MSCs were isolated from bone marrow of wild type (n=6) and *Sdc3^−/−^* (n=6) mice using a modified plastic adherence culture method^46^. Isolated murine MSCs were cultured in complete expansion medium (CEM) (Iscove Modified Dulbecco Medium (IMDM) (Gibco, UK) supplemented with 9% foetal bovine serum (FBS) (Gibco, UK), 9% horse serum (HS) (Gibco, UK) and 1% penicillin-streptomycin (Lonza, UK)) and used between passage 3 and 5. Cells were characterised in terms of their surface markers through immunophenotyping, growth kinetics by population doubling time (PDT) assay and colony forming units fibroblast (CFU-F) assay, and trilineage differentiation potential (adipogenic, osteogenic and chondrogenic) as previously described^17^. MSCs were tested for immunophenotype with the following antibodies: Ly-6A (Sca-1)-PE, CD44-PE, CD11b-PE, CD45-PE, CD105-PE, CD34-FITC and CD31-PE (all from eBioscience) using the FACSCanto II Flow Cytometer (BD Biosciences) and output data were assessed using Flowing Software 2.5.1 (Perttu Terho, Cell Imaging Core, Turku Centre for Biotechnology, University of Turku).

### 5.3 Adhesion assay

For cell adhesion assays, 96-well plates were coated with laminin, fibronectin or collagen type I at 1μg/cm^2^ (Sigma-Aldrich, UK) overnight at 4°C. After being blocked with BSA, 50 μl of 4×10^5^ cells/ml were plated to each well. Plates were incubated for 60 min, and non-adherent cells were washed with PBS, fixed with 4% formaldehyde and labelled with 5 mg/mL crystal violet for 10 min. Unbound dye was washed away with PBS, and the cells were lysed with 2% SDS before the absorbance at 550 nm was measured on FLUOstar OMEGA microplate reader (BMG Labtech, UK). Non-coated wells were also seeded and adhesion results were compared between the two cell genotypes. The percentage of cell attachment was determined relative to a set of standards with known cell numbers. Morphometric analysis was performed on 4% PFA-fixed, eosin (Sigma-Aldrich, UK)-stained cells. Images were acquired at 4x magnification using Olympus microscope IX-50 and quantitative data analysis was performed using ImageJ software (NIH, USA).

### 5.4 Migration: Scratch assay and transwell assay

For cell migration/scratch assays, 24-well plates were coated with 1 μg/cm^2^ fibronectin overnight at 4°C. 1×10^5^ cells were seeded and incubated at 37^0^C and 5% CO2 until 80% confluence. The cell monolayer was scratched in a straight line using a p200 pipet tip. Cells were washed with PBS to remove the debris and smoothen the edges of the scratch and then the culture medium was replaced with either CEM with full serum content (9% FBS and 9% HS, referred to as fCEM), CEM with reduced serum content (1% FBS and 1% HS, referred to as rCEM) or serum free medium (contains all components of CEM but neither FBS or HS, referred to as SFM) to prevent cell proliferation during migration. A digital image of each scratch was acquired using Olympus microscope IX-50. Cells were then incubated for 24 hours and all scratches digitally photographed again in the same location. For both time points ten vertical measurements of the cell layer gap were acquired, analysed using ImageJ software and the percentage of gap closure was calculated and compared between the two cell genotypes.

For transwell assays, 200 μl of 5×10^3^ cells in rCEM were seeded on the upper side of transwell inserts for 24-well plates (membrane 8.0μm pores) (Sarstedt, UK). 500 μl of serum (FBS and HS (1:1)), rCEM or 10 ng/ml of pleiotrophin in rCEM with 1% serum were added as chemoattractants in lower compartment. Serum free medium was used as control. Cells were incubated at 37°C and 5% CO2 for 15 h, fixed with 4 % paraformaldehyde for 15 min at RT and then washed with PBS. Cells were stained with DAPI for 10 min protected from light. Cells were counted from 15 random microscope fields for each sample using Olympus microscope IX-5. Quantitative data analysis was performed using ImageJ software and results presented as number of migrated cells.

### 5.5 Western blotting

Cells were lysed in RIPA buffer (R0278, Sigma) supplemented with protease and phosphatase inhibitor cocktails (Roche), then homogenised by passing the lysate through 21G needle and syringe. Lysates were clarified by centrifugation and total protein content analysed by BCA assay. For analysis of SDC3 protein levels, 100 ug of total protein was precipitated with methanol overnight at −20°C then centrifuged at 13,000 *x g* to pellet. Pellets were washed once with ice-cold acetone and resuspended in heparinase buffer (100 mM sodium acetate, 0.1 mM calcium acetate). Glycosaminoglycan chains were removed using 0.5 mU heparinase III (Ibex, Canada) and 0.5 mU chondroitinase ABC (Sigma) for 4 hours at 37 °C. Total protein 20 μg (or 40 μg for SDC3 analysis) was separated by SDS-PAGE and transferred onto nitrocellulose membranes (Hybond, GE Healthcare). Primary antibodies used were: p-AKT (1:3000, #4060, CST), AKT (1:3000, #9272, CST), p-ERK (1:3000, #9101, CST), ERK (1:3000, #9102, CST), p-Tyrosine (1:2000, #8954, CST), GAPDH (1:8000, G8795, Sigma) and SDC3 (0.14 μg/mL, AF2734, R&D). Secondary HRP-conjugated antibodies were used at 1:10,000 (Santa Cruz). Membranes were visualised using chemiluminescence (Clarity ECL, Bio-Rad) and imaged using ImageQuant-Las4000 (GE Healthcare). Band intensity was analysed using ImageJ.

### 5.6 Antigen-induced arthritis (AIA)

Animal procedures were undertaken in accordance with Home Office project licence P0F90DE46. AIA was induced in 7- to 8-week-old male C57Bl/6 mice (7–8 weeks) as previously described^47^. Briefly, mice were immunised subcutaneously with 1 mg/ml of methylated BSA (mBSA) emulsified with an equal volume of Freund’s complete adjuvant and injected intraperitoneally with 100 ml heat-inactivated Bordetella pertussis toxin, the subsequent immune response was boosted 1 week later. Then 3 weeks after the initial immunisation, AIA was induced by intra-articular injection of 10 mg/ml mBSA into the right knee joint and PBS into the left knee joint to serve as a control. Animals were inspected daily for arthritis development by measuring knee joint diameters using a digital micrometer (Kroeplin GmbH). The difference in joint diameter between the arthritic (right) and non-arthritic control (left) in each animal gave a quantitate measure of swelling (in mm).

### 5.7 Intra-articular injection of MSC

After one day post arthritis induction, 10 μl of SFM, containing 5×10^5^ wild type or *Sdc3^−/−^* MSCs were injected intra-articularly (0.5 mL monoject (29 G) insulin syringe, BD Micro-Fine, Franklyn Lakes, USA) through the patellar ligament into the right knee joint. Control animals were injected with SFM. Joint diameters were measured at 1, 2, 3, 5 and 7 days post injection. All measures were taken to reduce animal numbers (n = 4 per time point).

### 5.8 Arthritis index

Animals were sacrificed for histological analysis at days 3 and 7. Joints were fixed in 10% neutral buffered formal saline and processed as described previously^48^. H&E sections were scored for hyperplasia of the synovial intima (0 = normal to 3 = severe), cellular exudate (0 = normal to 3 = severe) and synovial infiltrate (0 = normal to 5 = severe); and Toluidine Blue stained sections for cartilage loss (0 = normal to 3 = severe) by two independent observers blinded to experimental groups ^47^. Scores were summated, producing a mean arthritis index.

### 5.9 Statistical analysis

Statistical analysis was carried out on GraphPad Prism 5.0. A one-way ANOVA followed by Tukey post hoc test was used for multiple comparisons and a two-tailed Student t test for comparisons of two variables. A p-value < 0.05 was considered significant.

## 7 Acknowledgments

We wish to acknowledge the help and support of staff at the Liverpool John Moores University Life Science Support Unit. We thank P. Evans and M. Pritchard, RJAH Orthopaedic Hospital, for their expertise in histology.

## 8 Author Contributions

FKJ performed signalling experiments, analysed the related data, prepared all figures and wrote the manuscript. AS performed the adhesion, migration and phenotyping experiments and analysed the related data. AGK provided support and technical skills for the experimental work. MH provided support for *in vivo* work and histological experiments. RM provided support for histological investigations and discussed and commented on the manuscript. NRF contributed to experimental design. AP designed the research study, analysed data, wrote and supervised the manuscript. OK conceived and designed the research study, performed *in vivo* experiments, analysed data, wrote and supervised the manuscript. All authors reviewed the manuscript before submission.

## 9 Funding

Funding was received from the UK EPSRC/MRC CDT in Regenerative Medicine (Keele University, Loughborough University, the University of Nottingham (EP/F500491/1), Oswestry Rheumatology Association and School of Medicine, Keele University. AP was funded by a Wellcome Trust ISSF. FKJ was funded by a BBSRC-DTP PhD studentship.

## 10 Conflict of Interest

The authors declare that the research was conducted in the absence of any commercial or financial relationships that could be construed as a potential conflict of interest.

## Notes

### Competing Interest Statement

The authors have declared no competing interest.

## References

1 Lipowsky, H. H. Relative shedding of glycosaminoglycans from the endothelial glycocalyx during inflammation and their contribution to stiffness of the glycocalyx. Biorheology 56, 191–205, doi:10.3233/BIR-190225 (2019).

2 Iozzo, R. V. & Schaefer, L. Proteoglycan form and function: A comprehensive nomenclature of proteoglycans. Matrix Biol 42, 11–55, doi:10.1016/j.matbio.2015.02.003 (2015).

3 Olguin, H. C. & Pisconti, A. Marking the tempo for myogenesis: Pax7 and the regulation of muscle stem cell fate decisions. J Cell Mol Med 16, 1013–1025, doi:10.1111/j.1582-4934.2011.01348.x (2012).

4 Choi, Y., Chung, H., Jung, H., Couchman, J. R. & Oh, E. S. Syndecans as cell surface receptors: Unique structure equates with functional diversity. Matrix Biol 30, 93–99, doi:10.1016/j.matbio.2010.10.006 (2011).

5 Hsueh, Y. P. & Sheng, M. Regulated expression and subcellular localization of syndecan heparan sulfate proteoglycans and the syndecan-binding protein CASK/LIN-2 during rat brain development. J Neurosci 19, 7415–7425 (1999).

6 Afratis, N. A. et al. Syndecans – key regulators of cell signaling and biological functions. FEBS J 284, 27–41, doi:10.1111/febs.13940 (2017).

7 Chernousov, M. A. & Carey, D. J. N-syndecan (syndecan 3) from neonatal rat brain binds basic fibroblast growth factor. J Biol Chem 268, 16810–16814 (1993).

8 Cornelison, D. D. et al. Essential and separable roles for Syndecan-3 and Syndecan-4 in skeletal muscle development and regeneration. Genes Dev 18, 2231–2236, doi:10.1101/gad.1214204 (2004).

9 Fuentealba, L., Carey, D. J. & Brandan, E. Antisense inhibition of syndecan-3 expression during skeletal muscle differentiation accelerates myogenesis through a basic fibroblast growth factor-dependent mechanism. J Biol Chem 274, 37876–37884, doi:10.1074/jbc.274.53.37876 (1999).

10 Pisconti, A., Cornelison, D. D., Olguin, H. C., Antwine, T. L. & Olwin, B. B. Syndecan-3 and Notch cooperate in regulating adult myogenesis. J Cell Biol 190, 427–441, doi:10.1083/jcb.201003081 (2010).

11 Fisher, M. C., Li, Y., Seghatoleslami, M. R., Dealy, C. N. & Kosher, R. A. Heparan sulfate proteoglycans including syndecan-3 modulate BMP activity during limb cartilage differentiation. Matrix Biol 25, 27–39, doi:10.1016/j.matbio.2005.07.008 (2006).

12 Rauvala, H. et al. Heparin-binding proteins HB-GAM (pleiotrophin) and amphoterin in the regulation of cell motility. Matrix Biol 19, 377–387, doi:10.1016/s0945-053x(00)00084-6 (2000).

13 Erdman, R., Stahl, R. C., Rothblum, K., Chernousov, M. A. & Carey, D. J. Schwann cell adhesion to a novel heparan sulfate binding site in the N-terminal domain of alpha 4 type V collagen is mediated by syndecan-3. J Biol Chem 277, 7619–7625, doi:10.1074/jbc.M111311200 (2002).

14 Pacifici, M. et al. Syndecan-3: a cell-surface heparan sulfate proteoglycan important for chondrocyte proliferation and function during limb skeletogenesis. J Bone Miner Metab 23, 191–199, doi:10.1007/s00774-004-0584-1 (2005).

15 Pisconti, A. et al. Loss of niche-satellite cell interactions in syndecan-3 null mice alters muscle progenitor cell homeostasis improving muscle regeneration. Skelet Muscle 6, 34, doi:10.1186/s13395-016-0104-8 (2016).

16 Patterson, A. M. et al. Induction of a CXCL8 binding site on endothelial syndecan-3 in rheumatoid synovium. Arthritis Rheum 52, 2331–2342, doi:10.1002/art.21222 (2005).

17 Kehoe, O., Cartwright, A., Askari, A., El Haj, A. J. & Middleton, J. Intra-articular injection of mesenchymal stem cells leads to reduced inflammation and cartilage damage in murine antigen-induced arthritis. J Transl Med 12, 157, doi:10.1186/1479-5876-12-157 (2014).

18 Eustace, A. D. et al. Soluble syndecan-3 binds chemokines, reduces leukocyte migration in vitro and ameliorates disease severity in models of rheumatoid arthritis. Arthritis Res Ther 21, 172, doi:10.1186/s13075-019-1939-2 (2019).

19 Dimarino, A. M., Caplan, A. I. & Bonfield, T. L. Mesenchymal stem cells in tissue repair. Front Immunol 4, 201, doi:10.3389/fimmu.2013.00201 (2013).

20 Lo Sicco, C. et al. Mesenchymal Stem Cell-Derived Extracellular Vesicles as Mediators of Anti-Inflammatory Effects: Endorsement of Macrophage Polarization. Stem Cells Transl Med 6, 1018–1028, doi:10.1002/sctm.16-0363 (2017).

21 Prockop, D. J. & Oh, J. Y. Mesenchymal stem/stromal cells (MSCs): role as guardians of inflammation. Mol Ther 20, 14–20, doi:10.1038/mt.2011.211 (2012).

22 Cahill, E. F., Tobin, L. M., Carty, F., Mahon, B. P. & English, K. Jagged-1 is required for the expansion of CD4+ CD25+ FoxP3+ regulatory T cells and tolerogenic dendritic cells by murine mesenchymal stromal cells. Stem Cell Res Ther 6, 19, doi:10.1186/s13287-015-0021-5 (2015).

23 Schofield, K. P., Gallagher, J. T. & David, G. Expression of proteoglycan core proteins in human bone marrow stroma. Biochem J 343 Pt 3, 663–668 (1999).

24 Beauvais, D. M., Burbach, B. J. & Rapraeger, A. C. The syndecan-1 ectodomain regulates alphavbeta3 integrin activity in human mammary carcinoma cells. J Cell Biol 167, 171–181, doi:10.1083/jcb.200404171 (2004).

25 McQuade, K. J., Beauvais, D. M., Burbach, B. J. & Rapraeger, A. C. Syndecan-1 regulates alphavbeta5 integrin activity in B82L fibroblasts. J Cell Sci 119, 2445–2456, doi:10.1242/jcs.02970 (2006).

26 Beauvais, D. M., Ell, B. J., McWhorter, A. R. & Rapraeger, A. C. Syndecan-1 regulates alphavbeta3 and alphavbeta5 integrin activation during angiogenesis and is blocked by synstatin, a novel peptide inhibitor. J Exp Med 206, 691–705, doi:10.1084/jem.20081278 (2009).

27 Beauvais, D. M. & Rapraeger, A. C. Syndecan-1 couples the insulin-like growth factor-1 receptor to inside-out integrin activation. J Cell Sci 123, 3796–3807, doi:10.1242/jcs.067645 (2010).

28 Hamaia, S. & Farndale, R. W. Integrin recognition motifs in the human collagens. Adv Exp Med Biol 819, 127–142, doi:10.1007/978-94-017-9153-3_9 (2014).

29 Leitinger, B. Transmembrane collagen receptors. Annu Rev Cell Dev Biol 27, 265–290, doi:10.1146/annurev-cellbio-092910-154013 (2011).

30 Heino, J. Cellular signaling by collagen-binding integrins. Adv Exp Med Biol 819, 143–155, doi:10.1007/978-94-017-9153-3_10 (2014).

31 Jones, G. W., Hill, D. G., Sime, K. & Williams, A. S. In Vivo Models for Inflammatory Arthritis. Methods Mol Biol 1725, 101–118, doi:10.1007/978-1-4939-7568-6_9 (2018).

32 Barbara, N. P., Wrana, J. L. & Letarte, M. Endoglin is an accessory protein that interacts with the signaling receptor complex of multiple members of the transforming growth factor-beta superfamily. J Biol Chem 274, 584–594, doi:10.1074/jbc.274.2.584 (1999).

33 Britten, M. B. et al. Infarct remodeling after intracoronary progenitor cell treatment in patients with acute myocardial infarction (TOPCARE-AMI): mechanistic insights from serial contrast-enhanced magnetic resonance imaging. Circulation 108, 2212–2218, doi:10.1161/01.CIR.0000095788.78169.AF (2003).

34 Jones, E. A. et al. Isolation and characterization of bone marrow multipotential mesenchymal progenitor cells. Arthritis Rheum 46, 3349–3360, doi:10.1002/art.10696 (2002).

35 Barnett, J. M., Suarez, S., McCollum, G. W. & Penn, J. S. Endoglin promotes angiogenesis in cell- and animal-based models of retinal neovascularization. Invest Ophthalmol Vis Sci 55, 6490–6498, doi:10.1167/iovs.14-14945 (2014).

36 De Rossi, G. & Whiteford, J. R. A novel role for syndecan-3 in angiogenesis. F1000Res 2, 270, doi:10.12688/f1000research.2-270.v1 (2013).

37 Somaiah, C. et al. Collagen Promotes Higher Adhesion, Survival and Proliferation of Mesenchymal Stem Cells. PLoS One 10, e0145068, doi:10.1371/journal.pone.0145068 (2015).

38 Vuoriluoto, K. et al. Syndecan-1 supports integrin alpha2beta1-mediated adhesion to collagen. Exp Cell Res 314, 3369–3381, doi:10.1016/j.yexcr.2008.07.005 (2008).

39 Hozumi, K., Suzuki, N., Nielsen, P. K., Nomizu, M. & Yamada, Y. Laminin alpha1 chain LG4 module promotes cell attachment through syndecans and cell spreading through integrin alpha2beta1. J Biol Chem 281, 32929–32940, doi:10.1074/jbc.M605708200 (2006).

40 Ogawa, T., Tsubota, Y., Hashimoto, J., Kariya, Y. & Miyazaki, K. The short arm of laminin gamma2 chain of laminin-5 (laminin-332) binds syndecan-1 and regulates cellular adhesion and migration by suppressing phosphorylation of integrin beta4 chain. Mol Biol Cell 18, 1621–1633, doi:10.1091/mbc.e06-09-0806 (2007).

41 Zwolanek, D. et al. beta1 Integrins Mediate Attachment of Mesenchymal Stem Cells to Cartilage Lesions. Biores Open Access 4, 39–53, doi:10.1089/biores.2014.0055 (2015).

42 Liang, C. C., Park, A. Y. & Guan, J. L. In vitro scratch assay: a convenient and inexpensive method for analysis of cell migration in vitro. Nat Protoc 2, 329–333, doi:10.1038/nprot.2007.30 (2007).

43 Yao, J., Zhang, L. L., Huang, X. M., Li, W. Y. & Gao, S. G. Pleiotrophin and N-syndecan promote perineural invasion and tumor progression in an orthotopic mouse model of pancreatic cancer. World J Gastroenterol 23, 3907–3914, doi:10.3748/wjg.v23.i21.3907 (2017).

44 Imai, S. et al. Osteoblast recruitment and bone formation enhanced by cell matrix-associated heparin-binding growth-associated molecule (HB-GAM). J Cell Biol 143, 1113–1128, doi:10.1083/jcb.143.4.1113 (1998).

45 Popov, C. et al. Integrins alpha2beta1 and alpha11beta1 regulate the survival of mesenchymal stem cells on collagen I. Cell Death Dis 2, e186, doi:10.1038/cddis.2011.71 (2011).

46 Peister, A. et al. Adult stem cells from bone marrow (MSCs) isolated from different strains of inbred mice vary in surface epitopes, rates of proliferation, and differentiation potential. Blood 103, 1662–1668, doi:10.1182/blood-2003-09-3070 (2004).

47 Nowell, M. A. et al. Soluble IL-6 receptor governs IL-6 activity in experimental arthritis: blockade of arthritis severity by soluble glycoprotein 130. J Immunol 171, 3202–3209, doi:10.4049/jimmunol.171.6.3202 (2003).

48 Kehoe, O. et al. Syndecan-3 is selectively pro-inflammatory in the joint and contributes to antigen-induced arthritis in mice. Arthritis Res Ther 16, R148, doi:10.1186/ar4610 (2014).

