## Supplementary figures and images for "MSCs from syndecan-3 null mice exhibit enhanced adhesion to collagen type I, hyperactivation of the AKT pathway and increased efficacy in inflammatory arthritis"

### Supplemental Figure 1

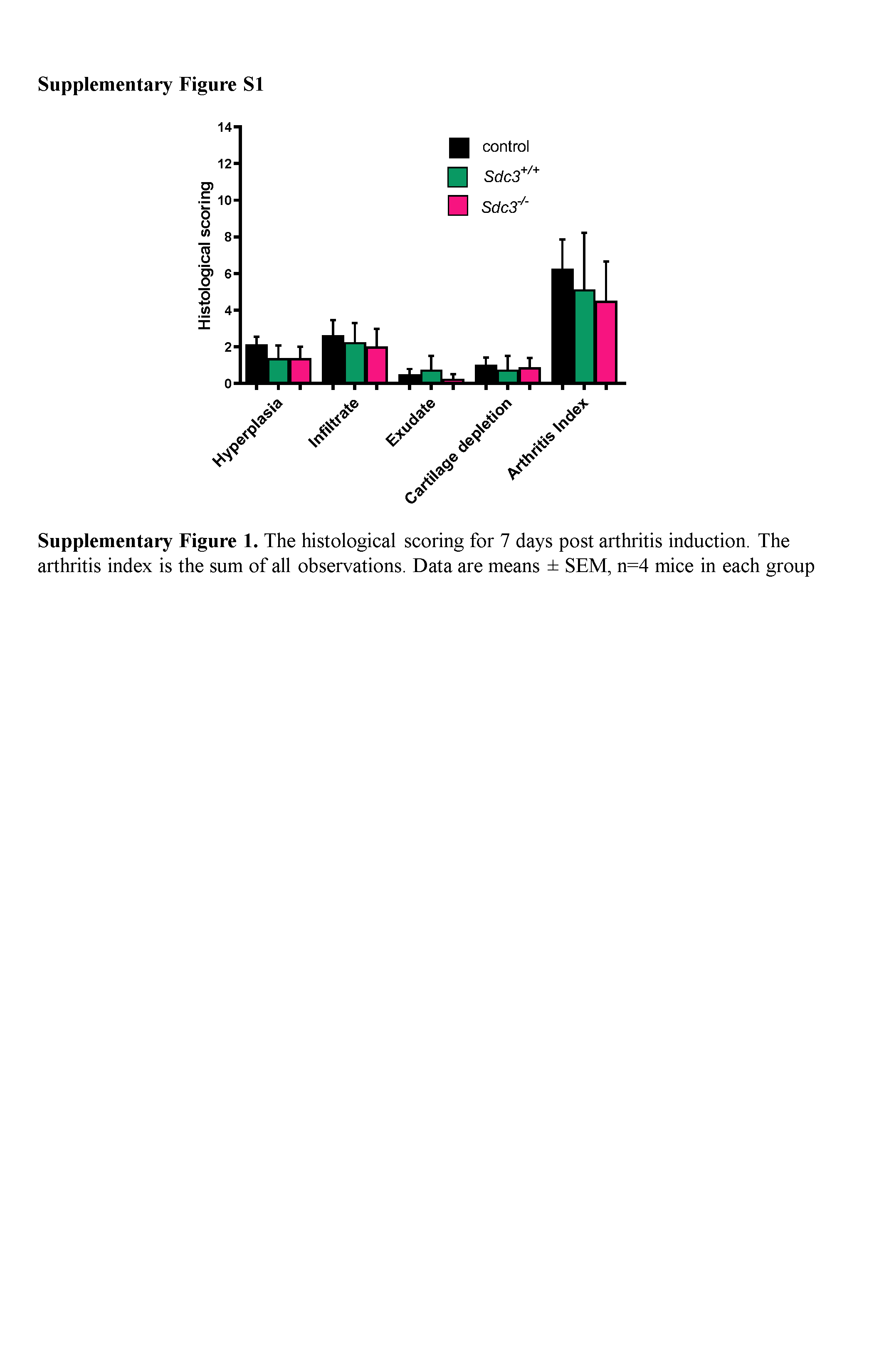
